# Human melanocyte development and melanoma dedifferentiation at single cell resolution

**DOI:** 10.1101/2020.05.25.115287

**Authors:** Rachel L. Belote, Daniel Le, Ashley Maynard, Ursula E. Lang, Adriane Sinclair, Vicente Planells-Palop, Laurence Baskin, Aaron D. Tward, Spyros Darmanis, Robert L. Judson-Torres

## Abstract

In humans, epidermal melanocytes are responsible for skin pigmentation, defense against ultraviolet radiation, and the deadliest common skin cancer, melanoma. While there is substantial overlap in melanocyte development pathways between different model organisms, species dependent differences are frequent and the conservation of these processes in human skin remains unresolved^1–3^. Thus, the biology of developing and adult human melanocytes remains largely uncharacterized. Here, we used a single-cell enrichment and RNA-sequencing pipeline to study human epidermal melanocytes derived directly from skin, capturing transcriptomes across different anatomic sites, developmental age, sexes, and multiple skin tones. Using donor-matched skin from distinct volar and non-volar anatomic locations, we uncovered subpopulations of melanocytes exhibiting site-specific enrichment that occurs during gestation and persists through adulthood. In addition, we identified human melanocyte differentiation transcriptional programs that are distinct from gene signatures generated from model systems. Finally, we use these programs to define patterns of dedifferentiation that are predictive of melanoma prognosis. Overall, the characterization of human melanocytes fresh from skin revealed new subpopulations, human-specific transcriptional programs, and valuable insights into melanoma dedifferentiation.

## INTRODUCTION

Epidermal melanocytes, the pigment producing cells of human skin, are responsible for skin tone and orchestrate the primary defense against damage from ultraviolet (UV) radiation. Some anatomic site-specific differences in pigmentation are due to environmental factors, such as the tanning response to UV exposure. Others, like the hypopigmentation at volar sites (such as palms and soles), are present at birth. In adult skin, mesenchymal – epithelial interactions are known to influence anatomic site-specific melanocyte survival and pigment production^4^ but melanocyte intrinsic factors that contribute to site-specific specialization remain unclear.

Model organisms are powerful tools for investigating melanocyte development. In chick and mouse, a transient, multipotent neural crest cell population gives rise to committed immature melanocyte precursors, called melanoblasts, via two spatially and temporally distinct pathways^2,3^. Such studies focus primarily on melanocytes in skin appendages (hair follicle, feather, and sweat gland). However, despite constituting the predominate subtype in human skin, resident epidermal melanocytes have not been the subject of analogous investigations into developmental trajectories and anatomic-specializations.

Melanocytes can give rise to melanomas which present distinct phenotypic and genomic characteristics correlated with primary tumor location^5,6^. Like many cancers, melanoma progression is coupled to dedifferentiation of the cell of origin^7^. The aggressive nature of melanoma is proposed to be rooted in unique attributes of the melanocytic linage^8^. Decoding the transcriptome of epidermal melanocytes across the human body during development and in aged skin would provide further insight into the precise origins of melanoma and the developmental programs reacquired during progression.

### Multi-site scRNA-seq of normal human melanocytes

We performed single cell RNA sequencing (scRNA-seq) on 34 healthy skin specimens across multiple anatomic locations (leg, arm, foreskin, palm and sole) from 22 donors aged 9.5 fetal weeks (f.w.) to 81 years (Fig. 1a, Supplementary Table 1) representing multiple skin tones and sexes. For each specimen, the epidermis was enzymatically removed from the dermis and dissociated into a single cell suspension. Since melanocytes comprise a small fraction of the total epidermal cell mass, FACS was used to increase the capture rate of CKIT+ melanocytes within the basal layer^9–11^ (Fig. 1a, Extended Data Fig. 1a,b, and methods). Sorted cells were then processed using the Smartseq2 single-cell RNA-seq protocol^12^. After quality control (Extended Data Fig. 1c-e, methods) and iterative Louvain clustering (see methods), differential expression was used to annotate 9,719 cells into the following cell-types: melanocytes, keratinocytes, eccrine sweat gland cells, and three immune cell populations (Fig. 1b,c and Extended Data Fig. 1f-i). Individual cells were then designated as cycling or non-cycling based on expression of established marker genes (Extended Data Fig. 2a-c)^13^. To further investigate heterogeneity within melanocytes, we performed Louvain clustering on melanocytes alone. The resulting 10 clusters did not correspond with skin tone, sex, or donor. Based on the expression of the top 5 differentially expressed genes (DEGs) for each cluster, clusters were binned into four groups (Extended Data Fig. 3a-d). These groupings corresponded with developmental age (adult, neonatal, and fetal) as well as a small group of fetal melanocytes expressing established melanocyte stem cell genes (Fig. 1d and Extended Data Fig. 3f,g). Interestingly, evaluation of donor skin anatomic location presented volar vs non-volar sites as a possible source of heterogeneity within the adult and fetal groups (Fig. 1e).

**Figure 1:**
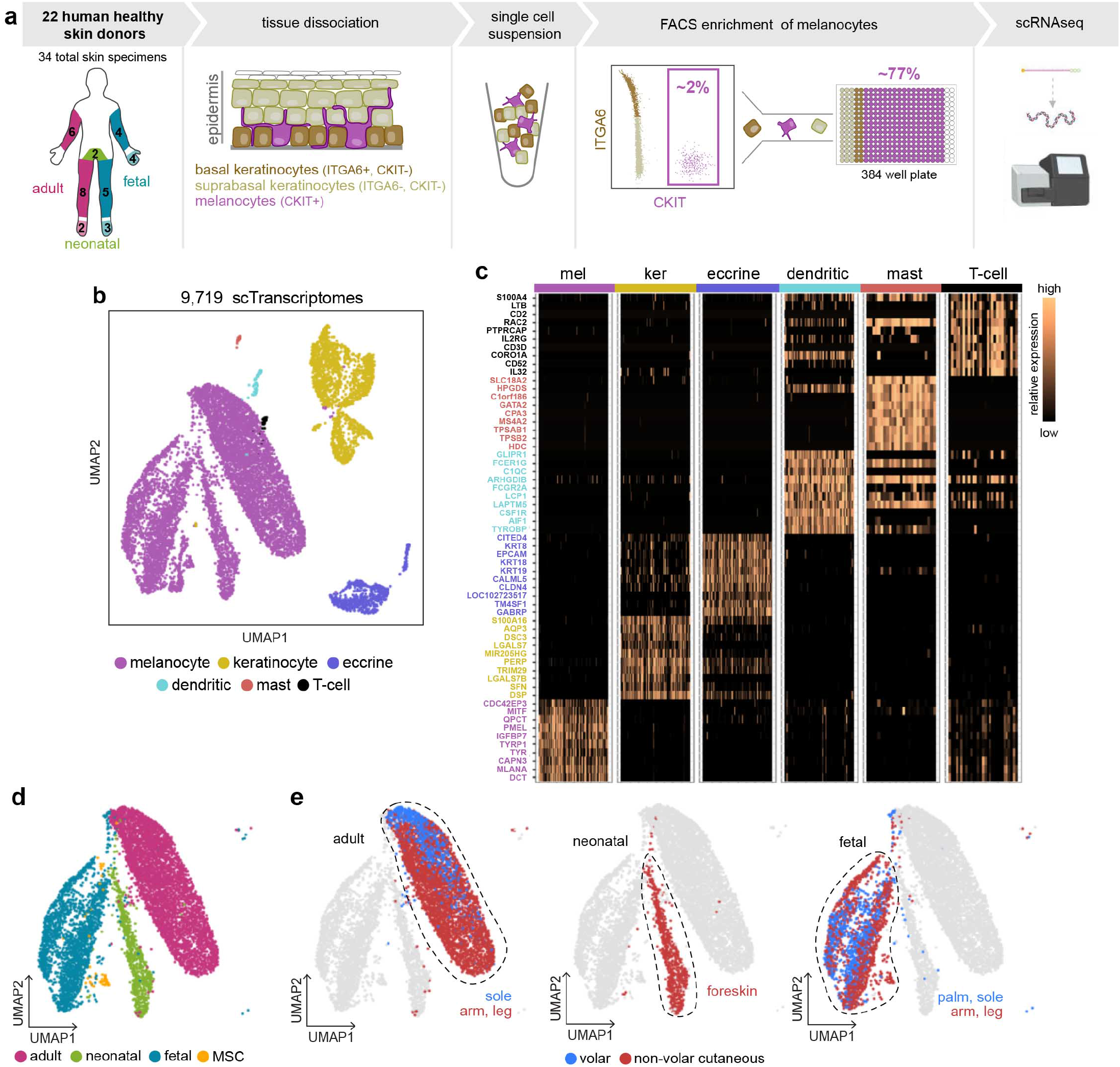
Melanocyte transcriptomic profiles differ based on development and anatomic location. **a)** Fresh from healthy human skin single cell isolation, enrichment, and sequencing pipeline. **b)** UMAP visualization of the 9,719 cells (7088 melanocytes, 1865 keratinocytes, 636 eccrine, 76 dendritic, 25 mast, and 29 T-cells) that passed quality control colored by cell types identified from Louvain clustering and candidate genes. **c)** Heat map showing the relative expression of top differentially expressed genes for each cluster in (b). **d)** UMAP of all non-cycling melanocytes with developmental age and fetal MSC annotation based on Louvain clustering (see also Extended Data Fig. 3). **e)** Fetal and adult melanocytes segregate by anatomic location.

### Anatomic site-enriched melanocyte sub-populations

The anatomical location of skin influences melanocyte survival and function but it remains unclear how site-specific specialization arises during melanocyte maturation^4^. UMAP segregation by anatomic location within the adult and fetal groups suggested melanocytes exhibit site-specific transcriptomic differences that arise early during development (Fig. 1e-f). To test this hypothesis, we queried donor matched volar and non-volar cutaneous specimens that spanned 10 f.w. to 77 years, different sexes, and different skin tones for transcriptional programs that distinguished volar vs non-volar cutaneous melanocytes across developmental ages (n=6 donors, n=17 skin specimens, Fig. 2a and Supplementary Table 1,*). Differential gene expression analysis (Mann-Whitney U test, Benjamini-Hochberg FDR < 5%) revealed 2,042 transcripts with site-specific expression in both fetal and adult donors (Fig. 2b-c, Supplementary Table 2). Using binary expression of the top 10 volar and non-volar cutaneous genes (see methods), we classified individual cells from the full cohort (n=22 donors, n= 34 skin specimens) as volar-like (v-mel) and non-volar cutaneous-like (c-mel). While v-mels and c-mels were present in all anatomic locations in both adult and fetal skin (Fig. 1d), v-mels were enriched in volar skin (mean: 94% ± 5% s.d. volar sites, ^~^7% ± 5 % non-volar sites) and c-mels were enriched in non-volar cutaneous skin (mean: ^~^89% ± 9% non-volar sites, 5% ± 5% volar sites). The presence of melanocytes with a c-mel signature in volar sites and melanocytes with the v-mel signature in cutaneous sites indicated: 1) two distinct sub-populations of epidermal melanocytes exist in human skin with anatomic site-specific enrichment, and 2) enrichment occurs during and persists after skin development. This discovery was validated via RNA FISH and immunofluorescence using the v-mel and c-mel signature genes that presented a striking level of inverse expression between volar and non-volar cutaneous melanocytes across all donor-matched skin: *NTRK2* (neurotrophic receptor tyrosine kinase 2) and *HPGD* (15-hydroxyprostaglandin dehydrogenase), respectively (Fig. 2e-h, Extended Data Fig. 4). These observations further suggest the previously reported site specific mesenchymal–melanocyte interactions^4,14^ that drive the epidermal phenotype in fully-developed skin, provide more permissive, but non-exclusive, conditions for one melanocyte subpopulation over another.

**Figure 2:**
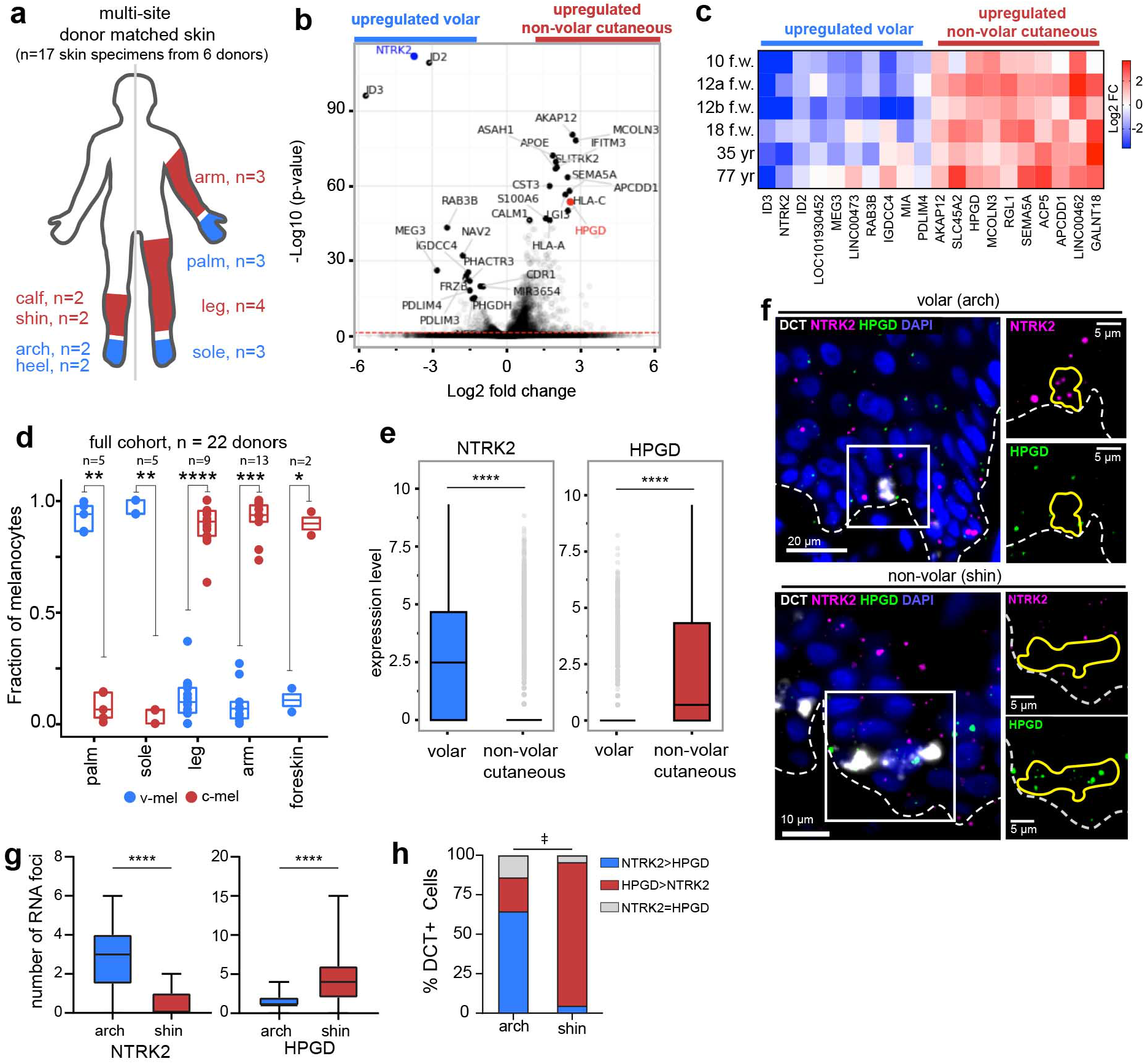
Anatomic site-specific melanocyte sub-population enrichment arises during development and persists in adulthood. **a)** Schematic illustrating cohort of donor-matched non-volar and volar skin. n = 6 donors and 17 total skin specimens. See also Extended Data Table 1. **b)** Volcano plot of genes enriched in non-volar cutaneous vs volar melanocytes. See also Supplementary Table 2. **c)** Top site-specific differentially expressed genes (DEGs), largest median per-patient log-fold-change between volar and non-volar melanocytes, among donor matched samples. **d)** Fraction of melanocytes with v-mel or c-mel signature in each skin specimen (n= 34) from all 22 patients. Two-sided Mann-Whitney U test with Bonferroni multiple testing correction,* p-value = 0.12, ** p-value <0.01, ***p-value < 0.001, ****p-value < 0.0001. **e)** Expression level of v-mel gene, *NTRK2*, and c-mel gene, *HPGD*, in volar melanocytes (sole, palm) compared to non-volar cutaneous melanocytes (arm, leg). Two-sided Mann-Whitney U test, ****p-value < 0.0001. Interquartile range with median, standard deviation, and outliers (grey circles). **f)** Representative pseudo-colored fluorescent microscopy images from RNAscope staining for *NTRK2, HPGD*, and the melanocyte marker *DCT* in adult volar and non-volar epidermis. Dashed line, epidermal-dermal melanocytes in volar (arch) and non-volar cutaneous skin (shin) in (f). Interquartile range, 5-95%, and outliers. Two-tailed unpaired t-test ****p-value < 0.0001 **h)** The percent v-mel (*NTRK2* > *HPGD*) and c-mel (*HPGD >NTRK2*) at each site in (f) showing volar skin has a significantly higher proportion of v-mels compared to non-volar cutaneous skin and non-volar cutaneous skin has a significantly higher proportion of c-mels compared to volar skin (‡, Two-sample Z-test for proportions, v-mel: z= 6.062, p-value = 1.8e-11, and c-mel: z= 7.885, p-value = 1.6e-15). For (g) and (h) n= 44 cells from arch and n= 22 cells from shin.

### Human-specific melanocyte differentiation programs

We captured three broad developmental ages in our cohort: fetal, neonatal, and adult. Consistent with the age based UMAP cluster arrangement, both diffusion pseudotime analysis and pairwise differential gene expression showed a progression from fetal to adult with neonatal melanocytes as an intermediate transcriptional state (Extended Data Fig. 5 and Supplementary Table 3). Our analyses did not reveal significant further transcriptional changes associated with donor age in adult melanocytes.

To identify gene signatures that best distinguished each human melanocyte developmental group, a regularized logistic regression model was trained using the single cell transcriptomes from 66% of the dataset representing the four developmental stages (Fig. 3a). The resultant <u>Dev</u>elopmental stage <u>Mel</u>anocyte (DevMel) model demonstrated excellent classification accuracy when tested on the holdout set, with f1-scores ranging from 0.93-1.00 (Extended Data Fig. 6a). Elastic net regularization yielded genes that collectively constituted developmental stage-specific expression programs: prg[MSC], prg[FET], prg[NEO] and prg[ADT] (Fig. 3b, Extended Data Fig. 6b-e and Supplementary Table 4).

**Figure 3:**
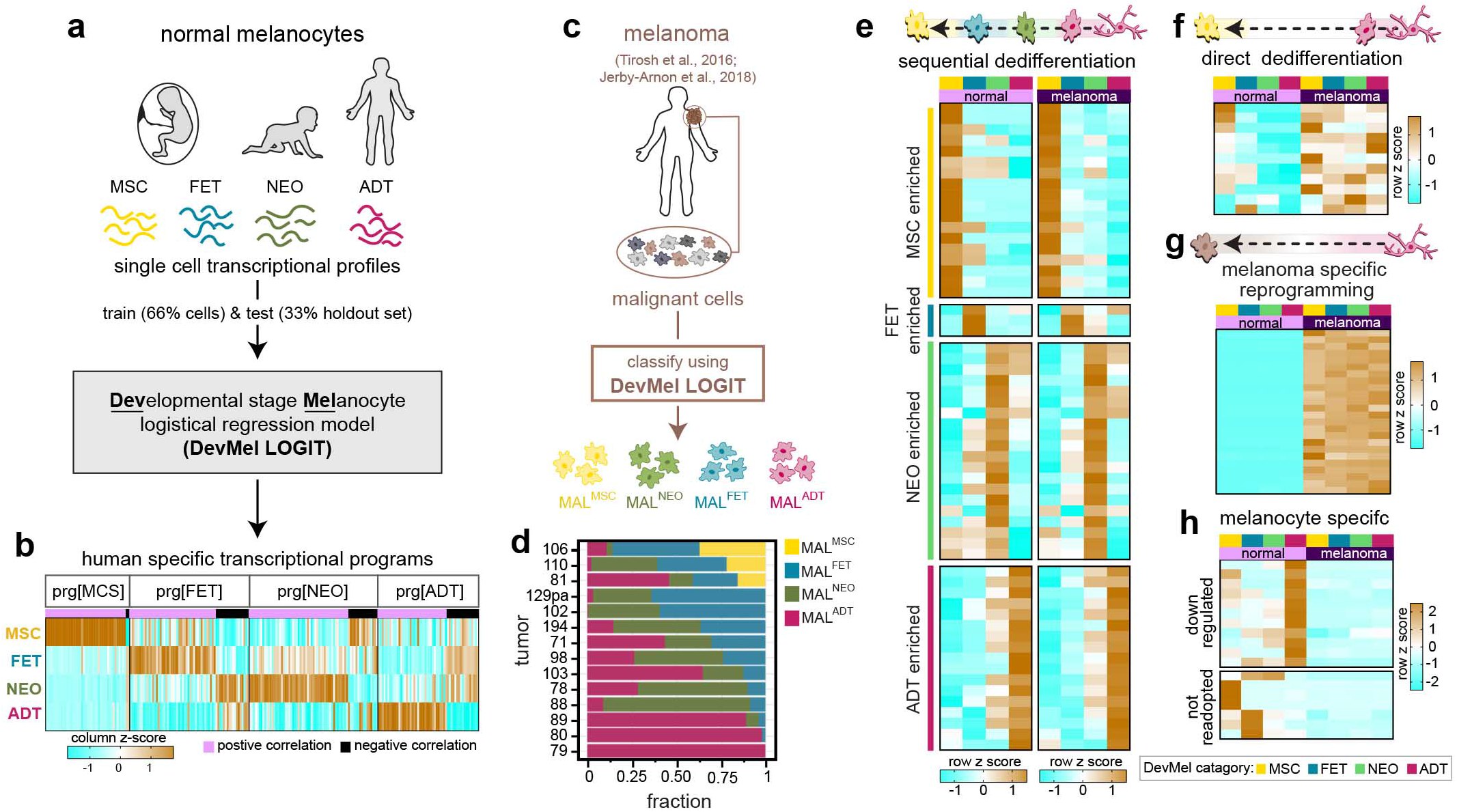
Identification of distinct patterns of developmental programs required in metastasized melanomas. **a)** Schematic of the Developmental stage Melanocyte logistical regression model (DevMel LOGIT) used to generate and validate unique transcription profiles for each developmental stage of normal human melanocytes. **b)** Heatmap of the relative expression (column z score) of genes in each DevMel program (prg) from (a). See also Extended Data Fig. 7 and Supplementary Table 4 for gene list. **c)** DevMel LOGIT was used to classify individual melanoma cells by normal melanocyte developmental stages. Every melanoma cell (MAL) was categorized by the predominantly expressed developmental stage program. **d)** Individual tumors are a heterogeneous mix of malignant cells in different dedifferentiation states. Fraction of MAL^ADT^, MAL^NEO^, MAL^FET^ and MAL^MSC^ cells in each of the 14 tumors analyzed from Tirosh *et al*. and Jerby-Arnon *et al*. in (c). **e-h**) Dedifferentiation can occur through several categories of cancer-associated transcriptional reprogramming: **e)** sequential dedifferentiation, a reverse stepwise unfolding of development; **f)** direct dedifferentiation, direct reacquisition of programs from early developmental stages; **g)** melanoma specific, acquisition of programs not associated with the stages of melanocyte development identified here. **h)** Normal adult developmental stage programs that are lost and earlier developmental stage programs not readopted in metastatic melanoma. Examples of each category are visualized as heatmaps of the relative expression (row z score). See Supplemental Table 5 for complete gene lists.

To benchmark human melanocyte development against known mammalian developmental systems, we assessed the expression of gene signatures previously defined during mouse melanocyte development^15–18^ and *in vitro* differentiation of human ES cells into mature melanocytes^19^ within adult (ADT), neonatal (NEO), fetal (FET) and melanocyte stem cells (MSC)s from c-mel enriched (non-volar cutaneous) skin (Extended Data Fig. 7 and Supplemental Discussion). Mouse melanocyte cell-type-specific signatures were more highly expressed in FET, NEO, and ADT melanocytes compared to MSC (p-value < 1e-12, Extended Data Fig. 7b,d). Alternatively, the mouse E14.5 and E17.5 melanoblast-specific signature^16^ was most highly expressed in MSC (> 1.5 fold change when compared to each other group, p-value < 1e-7, Extended Data Fig. 7c), suggesting the MSC cluster is associated with hair follicle development (see Supplemental Discussion). Consistent with this finding, the CD34+ mouse hair follicle melanocyte stem cell gene set^20^ was most highly expressed in MSC (p-value = 5.4e-38, Extended Data Fig. 7e). Of the *in vitro* differentiation programs, the mature differentiated melanocyte program was expressed across all developmental groups with the highest expression in FET, NEO and ADT groups compared to MSC (p-value < 1e-14, Extended Data Fig. 7f).

Comparison of DevMel programs to those identified in mouse or in human *in vitro* differentiation yielded sparse overlap (Extended Data Fig. 7g-j), indicating that our approach revealed previously unidentified programs specific to human fetal, neonatal, and adult skin. We therefore sought to determine whether profiles unique to human *in vivo* development could provide insight into melanoma dedifferentiation and aggression.

### Heterogenous reacquisition of developmental programs during tumorigenesis

Melanoma progression often coincides with the loss of melanocyte differentiation markers and upregulation of genes associated with earlier stages of development^16,20–23^. This process is broadly described as dedifferentiation. Given the known substantial cell-to-cell intratumor heterogeneity of melanoma^24^, we reasoned that single cells within a tumor might occupy various stages of dedifferentiation and that the proportion of cells in each state potentially influences overall patient outcome. To assess tumor heterogeneity as captured by the expression of human development-associated programs, we classified published single-cell malignant melanoma samples^25,26^ using our DevMel model (Fig. 3c). We observed inter- and intra-tumor heterogeneity in the representation of each melanoma group (Fig. 3d), indicating tumors are composed of a mix of dedifferentiated states in line with our initial hypothesis. Similar to our analysis of model system developmental programs, the MAL^MSC^ melanoma cells corresponded with stem cell-like transcriptional states (invasive, MITF-low, slow cycling) previously identified in human melanomas (Extended Data Fig. 8b-e, see Supplemental Discussion). In addition, we observed little developmental group specific enrichment for other reported melanoma signatures amongst the other MAL groups (Extended Data Fig. 8d). Taken together, these data showed classification of melanoma cells by human developmental programs generated a unique subdivision of non-stem like states, which could perhaps offer further insight toward understanding other states of differentiation in melanoma.

To better define the course of dedifferentiation during melanoma progression, we looked for differential gene expression patterns across each of the four MAL groups that were consistent with different forms of cellular reprogramming: (1) a retrograde unfolding of the differentiation cascade (sequential dedifferentiation)^20^, (2) direct reprogramming to a more pluripotent stage (direct dedifferentiation), or (3) the acquisition of a melanoma-specific program (Fig. 3b). Of 511 total unique genes, inclusive of DevMel model variables and MAL group top differentially expressed genes (Extended Data Fig.8a, Supplementary Table 5), 45% exhibited expression patterns consistent with sequential dedifferentiation, in which the relative expression across healthy melanocyte developmental groups was conserved among MAL groups (Fig. 3d). We found that 3.1% of genes exhibited a direct dedifferentiation pattern, indicating that expression of these genes may be a prerequisite for disease progression and metastasis (Fig. 3e). Supporting this interpretation, the small set of genes includes known markers of aggressive melanoma such as AXL and HMGA2^26,27^. Similarly, recently identified therapeutic resistance programs were evident in both the MSC healthy and MAL^MSC^ populations (Extended Data Figure 8e). We also identified genes expressed in healthy melanocyte groups that were down regulated in all melanoma groups (Fig. 3f), thus characterizing aspects of normal melanocyte expression that are either non-essential or potentially inhibitory to melanoma progression and/or metastasis.

Finally, 52 highly expressed genes in melanoma were absent from each of the healthy melanocyte developmental groups (Fig. 3g and Supplementary Table 5). Among the top differentially expressed genes was the melanoma-associated antigen PRAME, further supporting its use as a melanoma molecular diagnostic. Other of these “melanoma specific” genes might be important for melanoma progression or diagnosis such as the MTRNR2L family of transcripts, which encode for short peptides with anti-apoptotic activity^28^, and were highly and exclusively expressed in all melanoma groups.

### Developmental stage programs predict patient survival

To determine whether gene expression programs characteristic of different human developmental ages offer prognostic value, we applied CIBERSORT^29^ to estimate the fraction of melanoma cells similar to ADT, NEO, FET, MSC for all skin cutaneous melanoma (SKCM) tumor samples from The Cancer Genome Atlas (TCGA)^30^. Similar to the single cell melanoma dataset (Fig. 3d), we observed inter-tumor heterogeneity in the fractional representation of the four developmental groups (Fig. 4a). Hierarchical clustering of SKCM label distributions classified tumor samples according to the observed predominant developmental group: SKCM^ADT^, SKCM^NEO^, SKCM^FET^, SKCM^MSC^ (Supplementary Table 6). Neither genetic driver nor tumor site correlated with the developmental group classification of the tumor (Fig 4a; Extended Data Fig. 8f).

**Figure 4:**
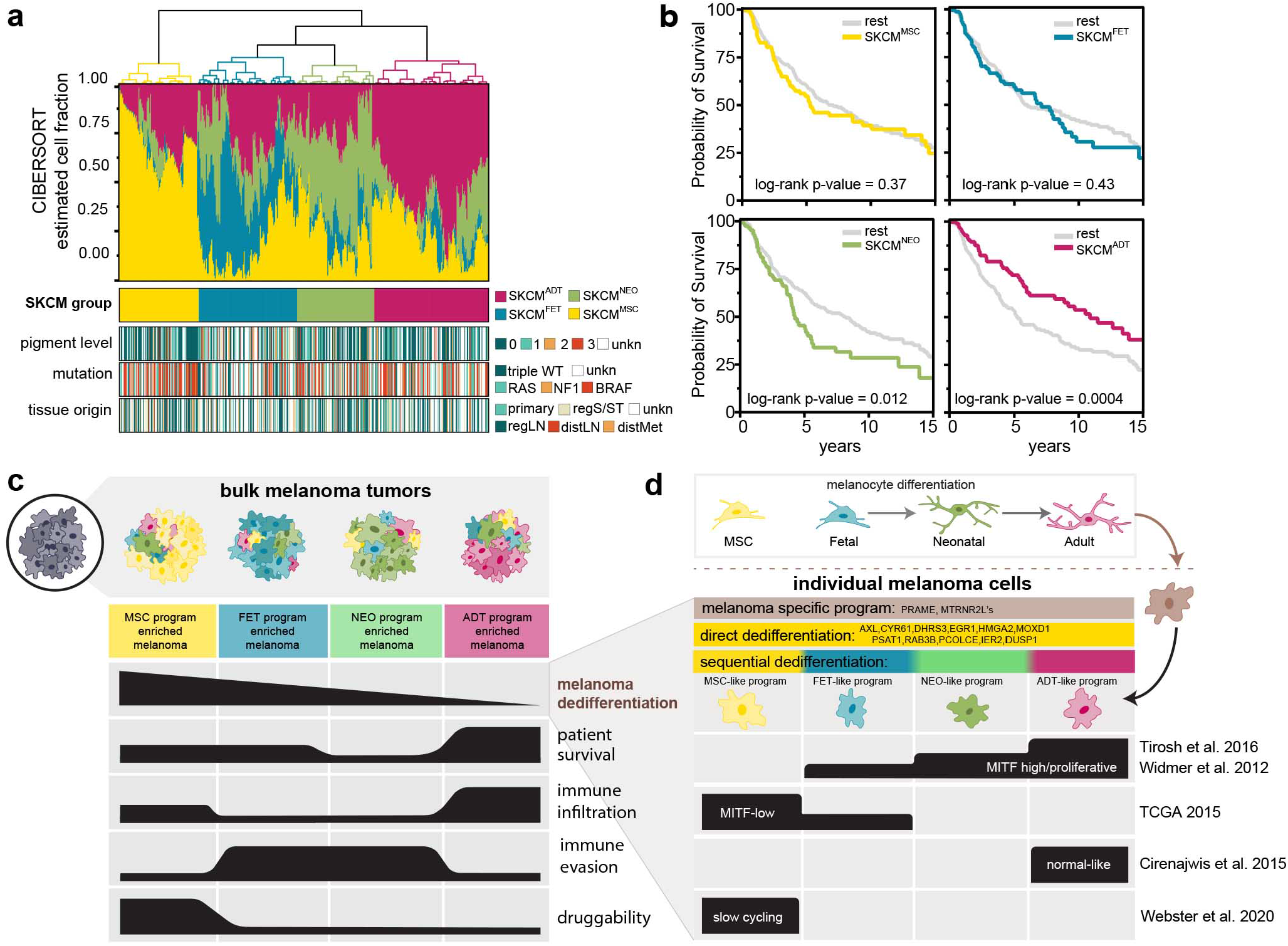
Reacquisition of specific developmental programs in heterogenous melanoma metastases is prognostic. **a**) Hierarchical clustering of TGCA SKCM tumors based on fractional composition of normal melanocyte developmental stages assigned using CIBERSORT (top) with clinicopathological features (bottom panels). **b)** Kaplan Meier curves for each SKCM group from (a). Enrichment for cells similar to ADT is associated with increased survival, whereas enrichment for NEO is associated with worse survival. **c-d)** Schematic summarizing the decoding of melanoma dedifferentiation using human developmental programs. **c)** Individual melanoma tumors are comprised of a heterogeneous mix of malignant cells expressing defined melanocyte developmental programs. The fraction of cells expressing each program within the tumor is predictive of overall survival and correlates to signatures of immune infiltration, evasion and potential therapeutic options. **d**) Each melanoma cell can occupy a different degree of dedifferentiation defined by sequential dedifferentiation transcriptional programs (Fig. 3 and Supplementary Table 5). The MSC- and adult-like programs are associated with previously described melanoma signatures whereas the fetal- and neonatal-like programs do not segregate with known melanoma signatures offering unique insight into previously uncharacterized melanoma transcriptional states (Extended Data Fig. 8). Genes common to all cells include melanoma specific genes, such as PRAME, or direct reprogramming to an early developmental stage (direct dedifferentiation).

Using our developmentally-defined subclasses of melanoma tumors, we evaluated the correlation of bulk tumor differentiation status with patient outcome. As expected, the most differentiated group (SKCM^ADT^) exhibited best median overall survival (SKCM^ADT^ = 11.0 yr vs rest = 5.3 yr). Surprisingly, the most dedifferentiated groups (SKCM^FET^, SKCM^MSC^) were not associated with worse outcomes (Fig. 4b); rather, the intermediately differentiated group (SKCM^NEO^) exhibited the shortest median overall survival (SKCM^NEO^ = 4.2 yr vs rest = 8.2 yr). To better understand this unexpected finding, we evaluated the expression of transcriptional programs associated with clinical response to therapeutics. Indeed, SKCM^NEO^ tumors expressed higher levels of transcripts associated with immune resistance^25^ (p-value=1.6e-2, SKCM^NEO^ vs rest) and a dearth in immune infiltration signatures^25^ (p-value=5.5e-4, SKCM^NEO^ vs rest) as well as FDA-approved therapeutic targets^31^ (p-value=1.6e-5, SKCM^NEO^ vs rest) (Extended Data Fig. 8g). The SKCM^MSC^ tumor group was unique in its increased expression of FDA-approved therapeutic targets (p-value=4.6e-21, SKCM^MSC^ vs rest). These data are in agreement with previous studies^20,25^ and demonstrate that while some amount of dedifferentiation is associated with worse prognosis, overall survival, immune evasion and immune resistance are not linearly correlated with dedifferentiation (Fig. 4c). Taken together, classification using human epidermal melanocyte developmental stage signatures revealed that at least four states of dedifferentiation constitute individual tumors, and that the proportion of melanocytes that have readopted a neonatal-like signature is associated with worse prognosis (Fig. 4c-d). The NEO program is not enriched for any previously identified melanoma transcriptional signature, suggesting its potential as prognostic biomarker remains untapped (Fig. 4d and Extended Data Fig. 8).

## DISCUSSION

We have provided a fresh-from-skin human epidermal melanocyte dataset that is, to our knowledge, the first human cell atlas entry that encompasses human development, sex and diverse race/ethnicities and includes multiple donor-matched anatomic locations.

We identified a novel population of epidermal melanocytes that appear early during human development. As the predominant class of melanocytes in volar regions, we speculate these v-mels could represent a distinct cell of origin of acral melanomas - a subtype of melanoma defined by its presentation in these regions and associated with distinct mutational landscapes and poor therapeutic response and overall survival^32,33^. It is possible epidermal v-mels are hypopigmented descendants of previously defined sweat gland stem cells^34^ and/or v-mels and c-mels are derived from the two known distinct lineage specification pathways^2^. Here we have provided markers that permit exploration of these hypotheses in future studies.

By characterizing melanoma dedifferentiation using human-specific developmental programs, our work sheds new light on the relationships among developmental stages, tumor characteristics, and melanoma cell transcriptional states (Fig. 4c,d). For example, with 63 years as the average age of melanoma diagnosis, our *in situ* adult melanocyte transcriptome provides a relevant basis for interrogating disease etiology and progression^35^. Moreover, our analyses identified the transcriptional state associated with neonatal melanocytes correlated to worst overall survival. Due to tissue availability and ease of culture, the neonatal melanocyte transcriptome is often considered the baseline “normal differentiated program” for comparison to melanoma transcriptomes. This technical artifact can explain why this program has been previously underappreciated. We further identified melanoma-specific genes directly acquired in all stages of dedifferentiation (Fig. 3f and Fig. 4d), suggesting that these genes may undergo positive selection during early metastatic dissemination. Along with the widely-accepted diagnostic melanoma biomarker PRAME^36^ and an established marker of invasion AXL^26^, we identified novel melanoma-associated genes. Further investigation into the mechanistic roles of these gene sets could reveal novel drivers of melanoma metastasis.

The findings presented here deliver a unique perspective on human melanocyte biology through the characterization of distinct transcriptional programs specific to development and function. Thus, the transcriptional programs identified here are valuable for understanding the diversity and malignant transformation of human melanocytes.

## ACKNOWLEDGMENTS

We thank the University of California, San Francisco Biospecimen Resource Program for their help with tissue acquisition, and Life Science Editors for critical editing of the manuscript. We thank the UCSF Program for Breakthrough Biomedical Research Sandler Fellows Program (to R.L.JT.) for financial support. We would like to thank Norma Neff and Michelle Tan for all the help with library quality control and sequencing.

## AUTHOR CONTRIBUTIONS

Conceptualization, R.L.B, D.L., A.D.T, S.D., and R.L.JT.; Methodology, R.L.B and A.M.; Validation, R.L.B. and D.L.; Formal Analysis, R.L.B. and D.L.; Investigation, R.L.B, D.L., and A.M; Resources, U.E.L., A.S., V.PP, L.B., and A.D.T.; Data Curation: D.L., and A.M.; Writing – Original Draft, R.L.B and R.L.JT.; Writing – Review & Editing, D.L., A.M., U.E.L. and S.D.; Visualization, R.L.B. and D.L.; Supervision, S.D. and R.L.JT.; Funding Acquisition, S.D. and R.L.JT.

---

**Extended Data Figure 1:**
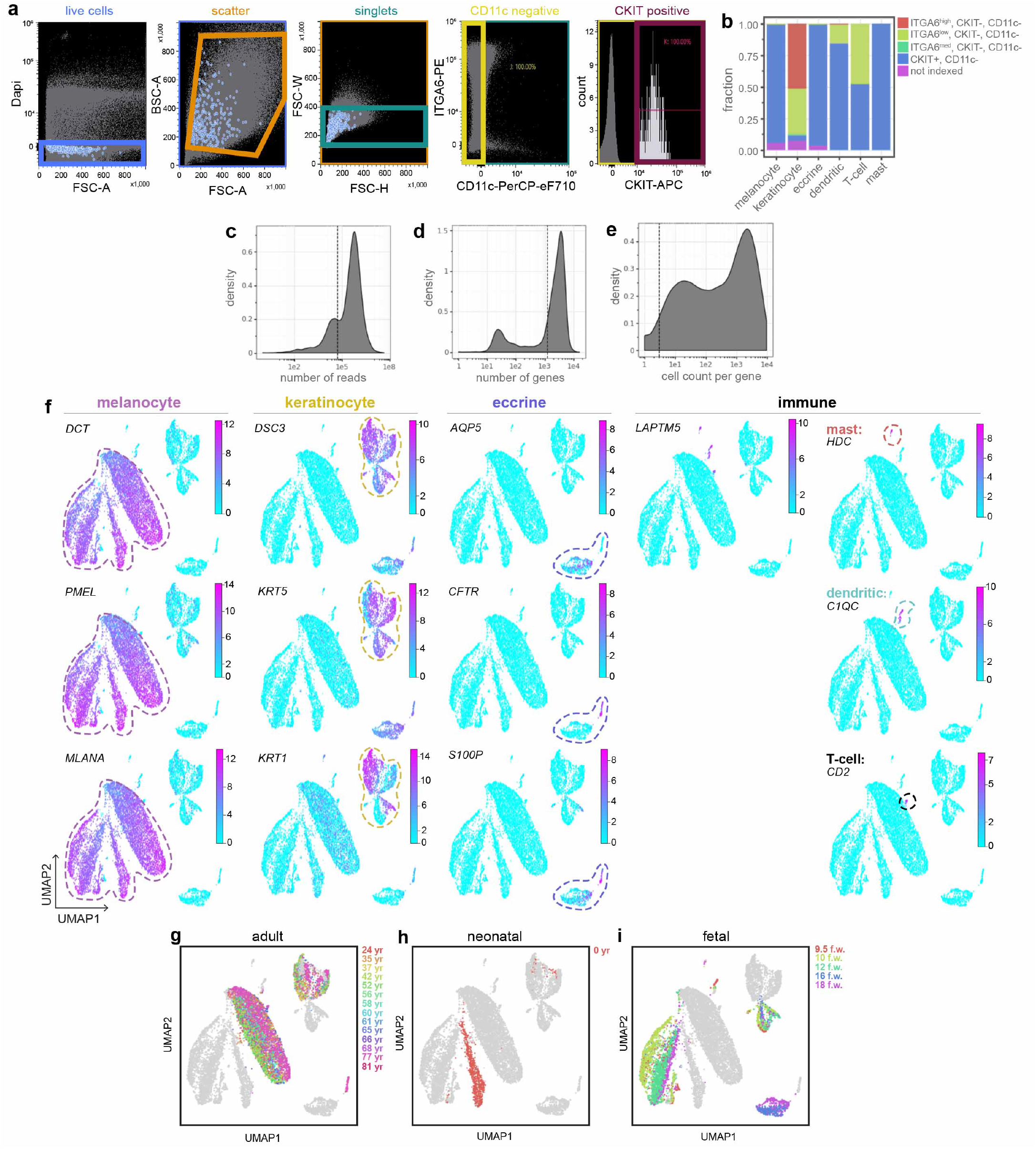
Single cell RNA sequencing quality control, cell-type specific markers, and donor age. **a)** FACS gate protocol for representative sort. Melanocytes (blue circles in live, scatter, and singlets) were isolated as CKIT+ cells from the CD11c-gate. **b)** Fraction of cells from each indexed FACS gate assignment. **c)** Number of reads and **d)** number of genes per cell for all 14,370 sequenced cells. Dashed line: quality control threshold, cells with < 50,000 reads and < 500 genes were excluded from further analysis. **e)** Genes expressed in more than 3 cells (dashed line) were included for subsequent analysis. **f)** Cell-type specific gene expression overlay on UMAPs. Genes indicated in upper left corner of each plot. **g-i)** UMAP with donor age overlay for **g)** adult, **h)** neonatal, and **i)** fetal cells.

**Extended Data Figure 2:**
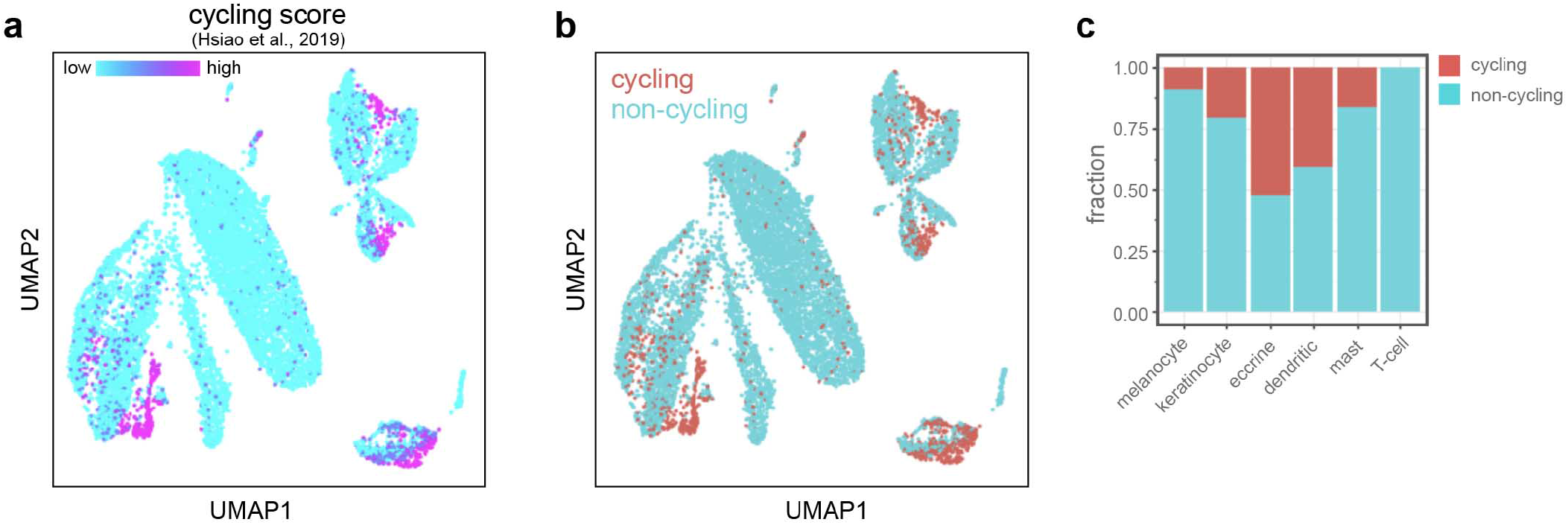
Characterization of cell cycle state. **a)** UMAP projection of cycling cell program score used to determine which cells were designated as **b)** cycling (blue, in G2 & M phase) vs non-cycling (red). **c)** Fraction of cycling and non-cycling cells for each cell type identified in Fig. 1.

**Extended Data Figure 3:**
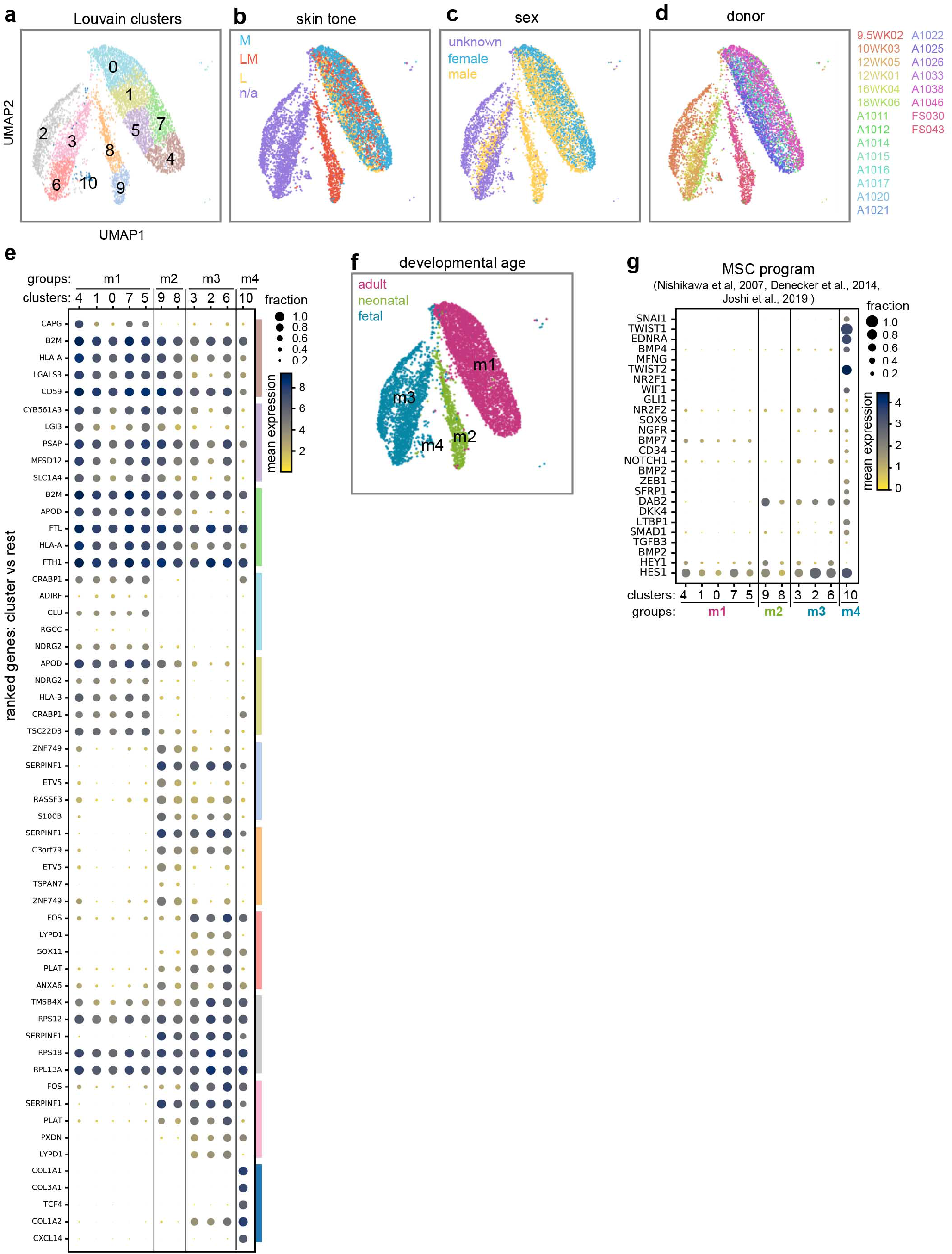
Identification and characterization of melanocyte clusters. **a)** UMAP of melanocytes showing Louvain clusters do not correspond to **b)** skin tone, **c)** sex, or **d)** donor. **e)** Dotplot of the mean expression and fraction of cells expressing the top 5 ranked genes (Wilcoxon Rank-sum) for each Louvain cluster in (a). Clusters with similar ranked gene expression patterns were bin into four groups: m1 (clusters 4,1,0,7,5), m2 (clusters 9,8), m3 (clusters 3,2,6), and m4 (cluster 10). **f)** Group m1 was comprised of adult melanocytes and m2 neonatal melanocytes, whereas groups m3 and m4 were comprised of fetal melanocytes. **g)** Dotplot showing group m4, from fetal hair-baring non-volar cutaneous skin (see Fig. 1e), expresses known melanocyte stem cell (MSC) markers. Groups are colored by developmental age in (f).

**Extended Data Figure 4:**
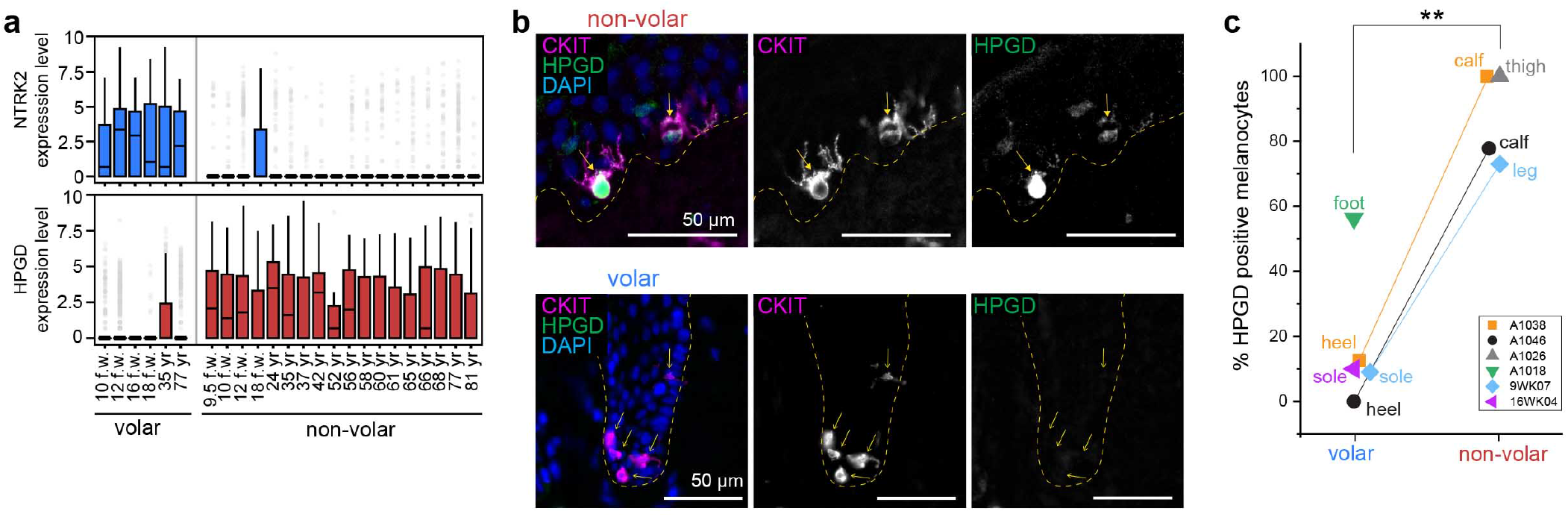
Validation of v-mel and c-mel site-specific enrichment in adult and fetal skin. **a)** Expression level of v-mel gene, *NTRK2*, and c-mel gene, *HPGD*, in volar melanocytes compared to non-volar cutaneous melanocytes at each age (n=22 donors). Interquartile range with median, standard deviation, and outliers (grey circles). **b)** Immunofluorescence co-staining of adult volar and non-volar skin cryo-sections with the c-mel marker HPGD (green) and melanocyte marker CKIT (magenta). Dashed line, epidermal-dermal junction. **c)** Quantification of site-specific enrichment of melanocyte c-mel and v-mel subclasses. Percent HPGD positive melanocytes per donor volar and non-volar skin. Adult skin: A1046, n=78 total cells; A1038, n = 39 total cells; A1018, n= 48 total cells; A1026, n= 15 total cells. Fetal skin: 9WK07, n= 41 total cells; 16WK04, n = 10 total cells. Two-tailed unpaired t-test, ** p-value = 0.001.

**Extended Data Figure 5:**
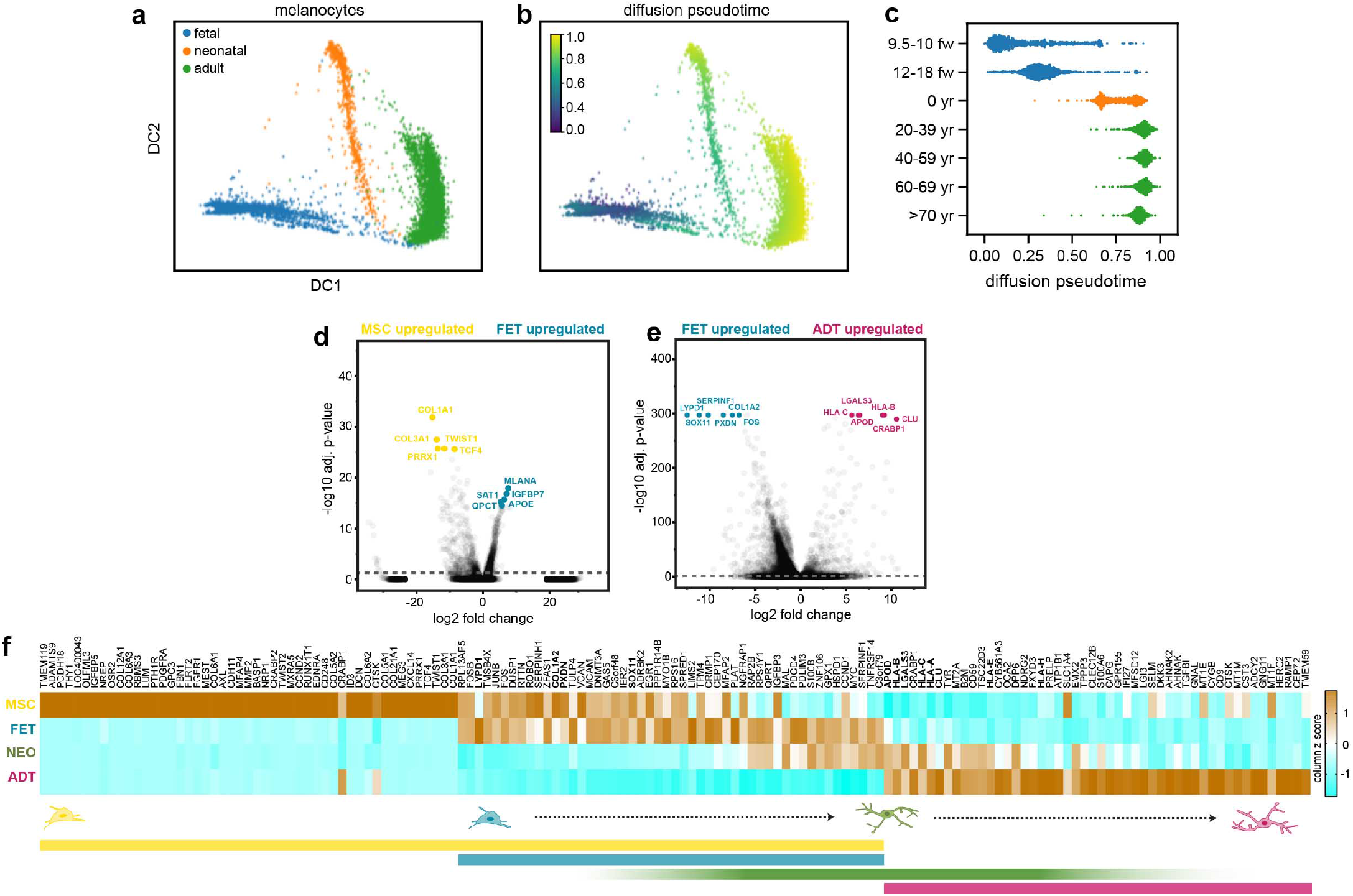
Pseudotime and pairwise differential expression analysis of developmental ages and groups. **a)** Melanocytes cluster by developmental age in diffusion component space DC1 and DC2. **b)** Pseudotime overlay onto DC space. **c)** Progression from fetal to adult through an intermediate neonatal transcriptional state. **d)** Volcano plot showing the top ten DEGs between MSC (yellow) and FET (teal) non-volar cutaneous melanocyte populations. **e)** Volcano plot showing the top ten DEGs between FET (teal) and ADT (magenta) non-volar cutaneous melanocytes. **f**) Heatmap visualization of the relative expression (column z score) of DEGs from (d) and (e) for all four non-volar cutaneous developmental groups. Both MSC and FET were enriched for known developmental genes (SOX11, LYPD1) and genes involved in extracellular matrix establishment/remodeling (COL1A2, PXDN). The ADT group expressed genes involved in innate immunity, inflammation and regulating apoptosis/cell stress in other cell types and tissues (HLAs, APOD, CLU, LGALS3). The NEO group exhibited high expression of a subset of genes from both the FET and ADT stages, consistent with neonatal melanocytes being an intermediate developmental state. See Supplementary Table 3 for the full list.

**Extended Data Figure 6:**
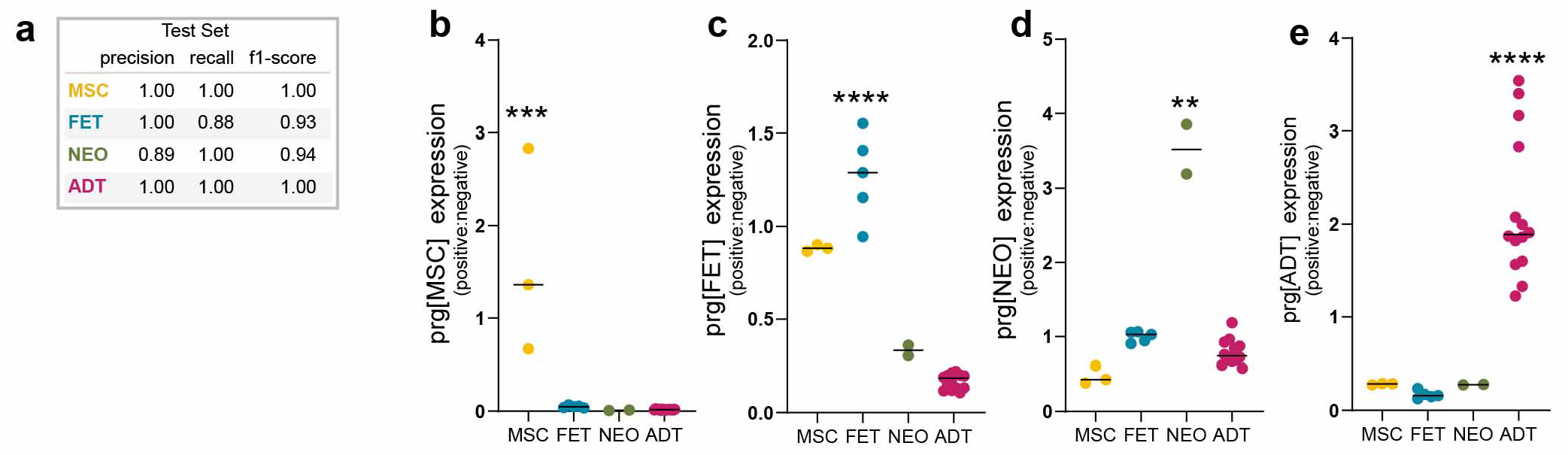
Transcriptional profiling of human melanocyte differentiation. **a)** DevMel LOGIT classifier performance on hold out validation set. **b-e)** DevMel program expression is highly expressed by cells from all skin donors within each corresponding developmental stage. Program expression for each donor (black line, average) is the ratio of the mean expression of positively correlated genes and negatively correlated genes. Significant by one-sided Mann Whitney U test. **b)** prg[MSC]: MSC vs rest *** p-value = 0.0005, **c)** prg[FET]: FET vs rest **** p-value 0.0001, **d)** prg[NEO]: FET vs rest ** p-value = 0.0072, **e)** prg[ADT]: ADT vs rest **** p-value < 0.0001.

**Extended Data Figure 7:**
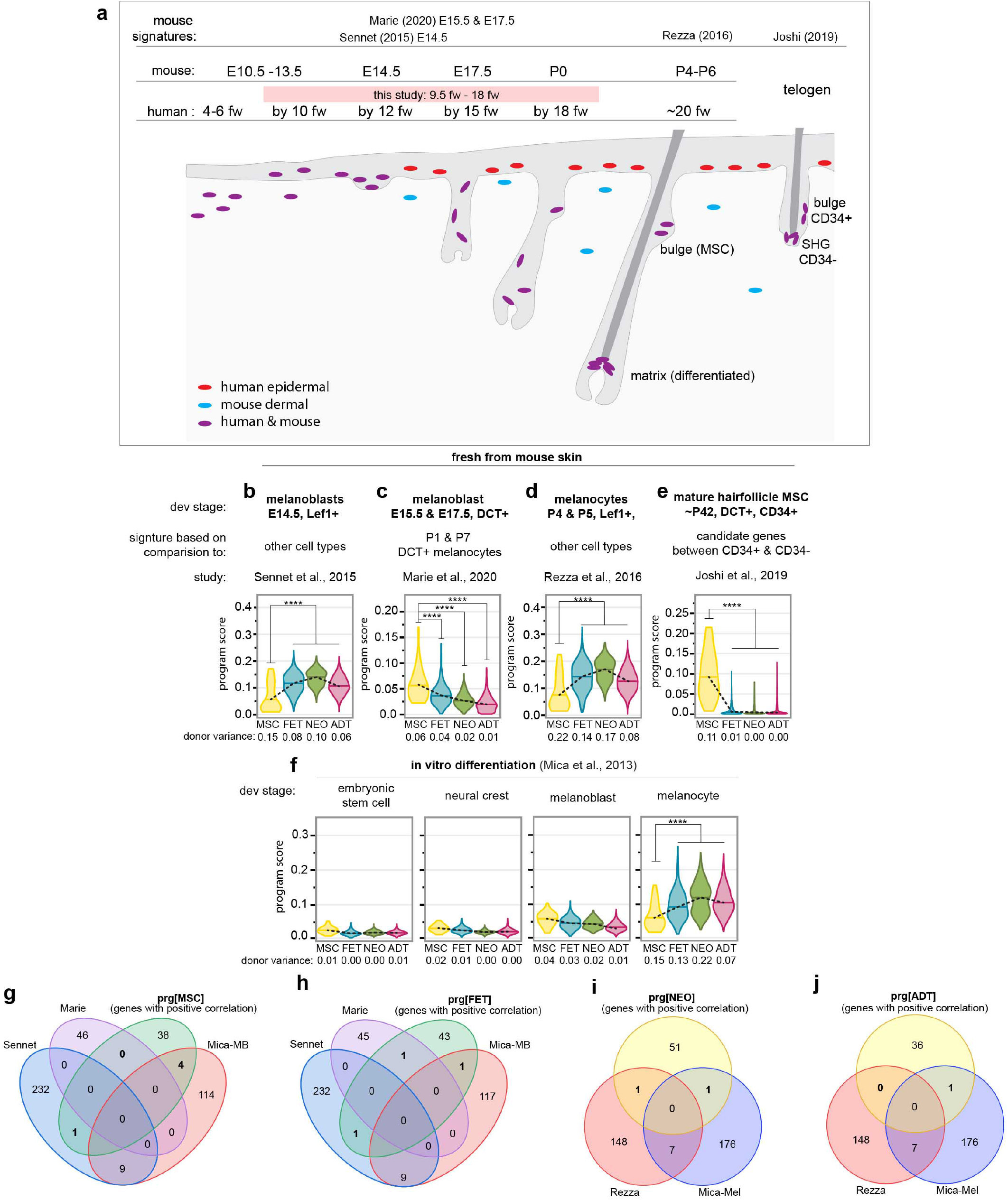
Evaluation of model mammalian melanocyte developmental program expression in human non-volar cutaneous melanocyte developmental groups. **a)** Schematic summarizing human^43,44^ and corresponding mouse^45^ melanocyte development captured in this study’s dataset. In hair-baring skin, both humans and mice develop follicular melanocytes (purple). Mice retain a dermal melanocyte population (blue) in fully developed skin, whereas humans develop resident epidermal melanocytes (red) within the skin at all anatomic locations. **b-f)** Violin plots show distribution of indicated transcriptional program expression scores for individual cells within each developmental group MSC, FET, NEO, and ADT. Dashed line: mean expression. Program scores were generated from published signatures of **b-c)** mouse melanoblasts (cell committed to the melanocyte fate)^16,18^, **d)** melanocytes^17^, and **e)** melanocyte stem cells from mature hair follicles in adult mice^15^ and **f)** *in vitro* melanocyte differentiation stages. For each individual cell, the program score is the mean normalized expression for all genes in the indicated published gene signature. The variance of the average program expression among donors within the MSC, FET, NEO, and ADT groups (reported below the corresponding group for each violin plot) was low showing concordance across ages within each group. **** p-value < 1 x 10^−7^, significance determined by one-sided Man-Whitney U test. **g-h)** Venn diagrams showing the number of unique and overlapping genes of melanoblast related gene signatures with the positive correlated component of the DevMel profiles **g)** prg[MSC] and **h)** prg[FET]. **i-j)** Venn diagrams showing the number of unique and overlapping genes of differentiated melanocyte related gene signatures with the positively correlated component of the DevMel profiles **i)** prg[NEO] and **j)** prg[ADT].

**Extended Data Figure 8:**
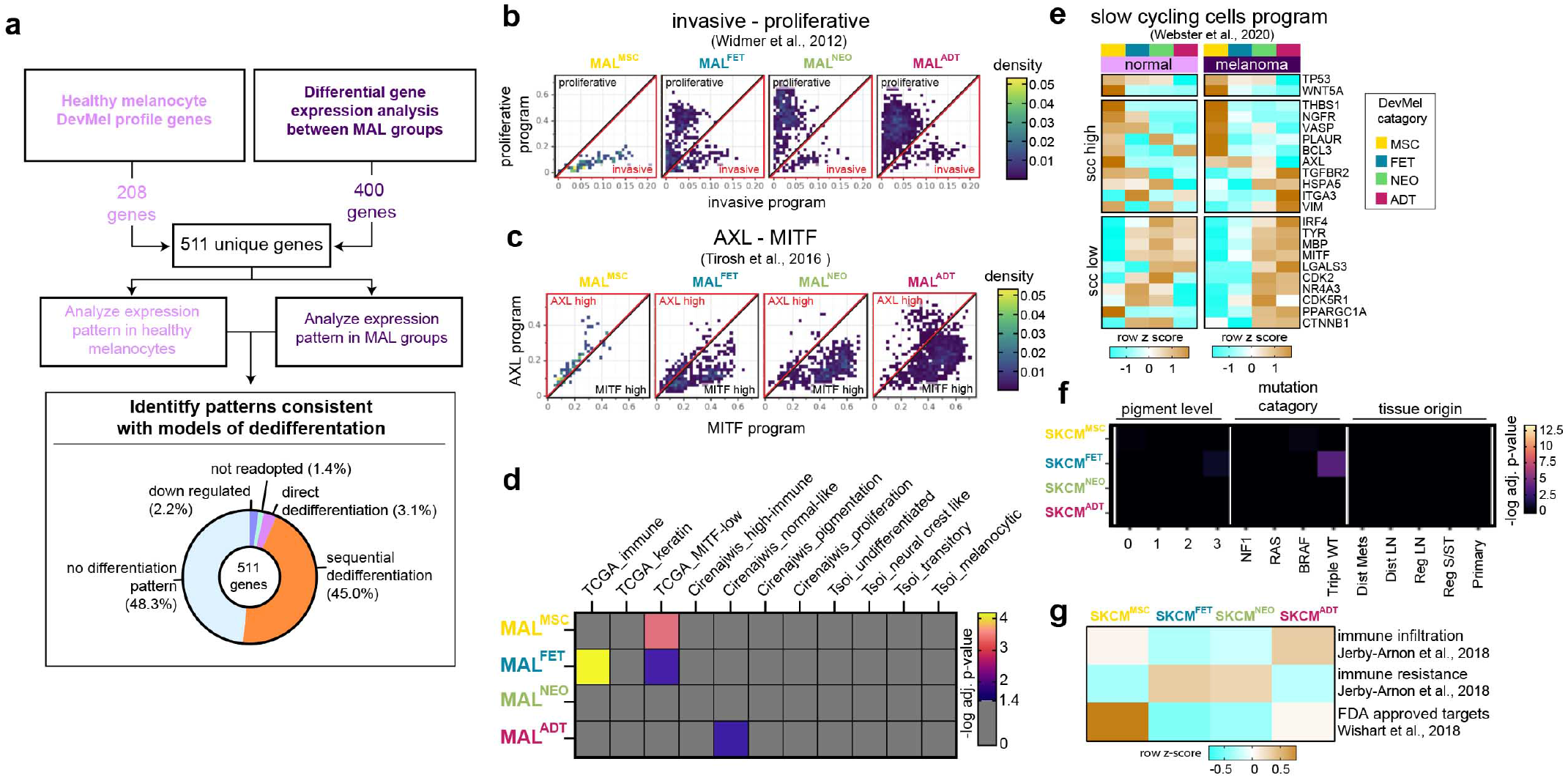
Characterization of melanoma cells and tumors classified by *in situ* human melanocyte developmental programs. **a)** Workflow to generate gene set (511 unique genes) used to identify patterns associated with melanoma dedifferentiation (top). Percent of genes across MAL groups that exhibit patterns consistent with dedifferentiation categories in Fig. 3e-g. **b-c)** Density plots showing the expression of **b)** the Widmer *et al*. invasive and proliferative programs and **c)** the Tirosh *et al*. AXL and MITF programs for individual cells in MAL^ADT^, MAL^NEO^, MAL^FET^ and MAL^MSC^ groups. **d)** Pairwise Fisher exact test showing negative log10 adjusted p-values for the gene set enrichment analysis conducted for TCGA *et al*., 2015, Cirenajwis *et al*., 2015 and Tsoi *et al*., 2018 gene signatures. **e)** Heatmap showing the relative expression levels (row z score) of WNT5A high, TP53 high slow cycling cell associated genes in each normal melanocyte and MAL developmental group. **f)** Pairwise Fisher exact test showing negative log10 adjusted p-values for clinicopathological feature and transcriptional categorization within each SKCM group (SKCM^ADT^, SKCM^NEO^, SKCM^FET^, SKCM^MSC^). There is little to no difference in the enrichment of pigment level, mutation category, or tissue origin between SKCM groups in Figure 4. **g)** Heatmap showing the relative expression levels (row z-score) of immune infiltration program, immune evasion program and FDA-approved therapeutic targets in SKCM groups.

## SUPPLEMENTARY INFORMATION

### SUPPLEMENTAL DISSCUSSION

#### Supplemental discussion related to Extended Data Figure 3

The single cell isolation procedure for adult skin was optimized for CKIT+ interfollicular epidermal melanocytes; thus, we expected exclusion of CKIT negative/low follicular melanocyte stem cells (MSC) (see methods). We therefore assessed the expression of known MSC genes^15,46,47^ in each cluster. The lack of expression for known follicular melanocyte stem cell (MSC) genes in group m1 (Extended Data Fig. 3g) suggests the strategy was successful in the adult skin. While, it is possible that CKIT+ differentiated follicular melanocytes were captured, they were not transcriptionally distinct from epidermal melanocytes. In contrast, two of the top 5 ranked genes for group m4, TC4F^48^ and CXCL14^49^ are associated with stem- and progenitor-cell function, suggesting MSCs were captured from fetal hair-baring, non-volar cutaneous skin (Fig. 1e and Supplemental Table 1, leg and arm). Consistent with this hypothesis, the putative MSC group expressed many of the known MSC genes unlike the other fetal, neonatal and adult clusters (Extended Data Fig. 3g).

#### Supplemental discussion related to Extended Data Figure 6

Melanocyte development has primarily been studied using non-human model organisms^3^ or employing *in vitro* differentiation of human pluripotent stem cells^19^. Gene signatures that define stages of melanocyte differentiation have been identified in these systems, including mouse CD34+ follicular melanocyte stem cells and melanoblasts^15,16^ and *in vitro* human embryonic stem cells, neural crest cells, melanoblasts, and differentiated melanocytes^50^. “Melanoblast” is a designation used to describe committed immature melanocytes along a broad range of developmental time points - from initial commitment to melanocyte fate, which occurs prior to epidermal infiltration through hair follicle morphogenesis, and along two distinct lineage specification pathways^3^. Given the fetal ages sampled here, we expected observation of *in vivo* human melanoblasts (Extended Data Fig. 7a). If the transcriptional programs of developing melanocytes were conserved from the model systems to *in vivo* human development, we would have expected substantial enrichment for the model system gene sets in the corresponding developmental stage. It was therefore surprising that the published mouse CD34+ melanocyte stem cell signature was the only signature to have relatively high and unique expression within the corresponding human developmental stage profiled here. The observed discordance could have several sources: species-, condition-, or isolation method-specific differences, differences between scRNAseq and bulk mRNAseq, differences in timescale of skin development between species, and/or the type of analysis used to generate the final gene signature.

Both the Sennet *et al*. E14.5 melanoblast signature (Extended Data Fig.7b) and the Rezza *et al*. P4 & P5 melanocyte signatures (Extended Data Fig.7d) were more highly expressed in the FET, NEO and ADT melanocytes compared to MSCs. In these studies, melanocytic cells were isolated using LEF1 and CKIT expression, and the gene signatures were derived from the comparison of melanocytic cells to other skin cells. In mice, LEF1 is a marker of differentiated (and differentiating) melanocytes and is not expressed in MSCs^51,52^. Thus, the resulting gene signatures represent a general melanocytic cell-type specific program, exclusive of MSCs, at each mouse developmental time point. The observed low program expression in the human MSC group defined in the current study is therefore consistent with the experimental design for the Sennet and Rezza studies (Extended Data Fig.7b,d). In contrast, the Marie et al. melanoblast signature (Extended Data Fig.7c) was derived from the comparison of DCT+ melanoblasts at E15.5 & E17.5 to P1 & P7 melanocytes, and is therefore a melanoblast specific signature. DCT is expressed in differentiated (and differentiating) melanocytes as well as MSCs. Unlike the Sennet et al. melanoblast signature, the Marie *et al*. melanoblast signature had decreased expression within more differentiated developmental ages, which is to be expected (Extended Data Fig. 7c). Interestingly, the MSC group had the highest expression. Mouse hair follicle morphogenesis occurs around E14 and is completed postnatally by P8 as a fully mature hair-bearing follicle in anagen phase^45^. In humans, hair follicle formation is reported to start around 10 f.w. with mature hair follicles appearing around 20 f.w. depending on anatomic location and study^43,44^. The fetal skin specimens in our dataset coincide with the onset and later stages of human hair follicle development, which would encompass morphological stages that resemble mouse hair follicle development at E15.5 and E17.5 from Marie *et al*. Unlike mice, human hair-baring skin contains both hair follicle-associated and epidermal-associated melanocytes. Therefore it is reasonable that the mouse melanoblast-specific program from Marie *et al*. is most highly expressed in a subset of the human fetal melanocytes that express known follicular-associated gene signatures (Extended Data Fig.3g and 7c,e).

The *in vitro* derived mature melanocyte gene set was most highly expressed in all groups, with enrichment among FET, NEO, and ADT melanocytes (Extended Data Fig7f). These observations suggest that *in vitro* generation of melanocytes from pluripotent stem cells does not distinguish between differentiating, young and aged melanocytes. Differentiation protocols that better distinguish the *in vivo* profiles reported here, especially accounting for the effect of the aged adult developmental state, would be a valuable tool for the field.

#### Supplemental Discussion Related to Extended Data Fig. 8

A number of studies have identified transcriptional programs associated with a variety of melanoma phenotypes and patient outcomes. Here, we identified four malignant melanoma states identified through classification by the most similar human developmental stage (the MAL groups: MAL^MSC^, MAL^FET^, MAL^NEO^, MAL^ADT^). We were interested in determining whether any of the MAL group signatures overlapped with, or were enriched for, previously published transcriptional signatures. Two well established transcriptional signatures have been defined as “proliferative / MITF high” and “invasive / AXL high”^26,53^. These classifications center on the average expression level of a MITF regulated transcriptional program (inclusive of MITF itself). Since increasing expression of the MITF program is thought to drive a progressively differentiated state^54^, we reasoned the MAL groups, derived from developmental signatures, might likewise present differential expression of the this program. In agreement with this model, the proportion of cells expressing the proliferative and MITF high programs also increased with more differentiated groups (MAL^ADT^ and MAL^NEO^) as compared to the less differentiated groups (Extended Data Fig. 8b-c). In particular, the MAL^MSC^ group comprised almost entirely of invasive melanoma cells and was devoid of MITF high cells, consistent with a dedifferentiated state (Extended Data Fig. 8b-c). Corroborating this interpretation, MAL^MSC^ cells were enriched for TCGA MITF-low signature genes and expressed higher levels of a gene signature indicative of a slow cycling cell therapeutic resistance state associated with stem cells (Extended Data Fig. 8d,e). Thus, we conclude that the stem-like melanoma cells identified here (MAL^MSC^) are akin to stem-like melanoma cells identified in previous studies. Interestingly, only 1 of the 14 tumors analyzed here was a treatment-naïve tumor (81) but it too contained a small MAL^MSC^ population (approximately 11% of tumor cells), consistent with previous reports describing the presence of a stem-like dedifferentiation state prior to therapy (Extended Data Fig. 8a). We further demonstrate that this program is also expressed in healthy melanocytes (the MSC cells), including enrichment for the slow cycling cell state (Extended Data Fig. 8e) implicating the transcriptional program associated with therapy-resistant, stress-induced, slow cycling tumor cells as a pre-existing feature of MSCs from normal skin as opposed to a feature solely acquired during tumorigenesis or therapeutic exposure.

Previous reports have used transcriptional signatures to classify cohorts of melanomas. Most notably, the TCGA cohort^55^ can be classified as “immune”, “keratin”, or “MITF-low”; and the Cirenajwis *et al*. cohort^56^ as “Immune”, “Normal-like”, “Pigmented” or “Proliferative”. Based upon profiling of *in vitro* differentiation of human stem cells into melanocytes, signatures for “undifferentiated”, “neural crest like”, “transition”, and “melanocytic” have also been reported^20^. We reasoned that the four malignant melanoma states identified here might correspond to the three or four classification signatures reported in these aforementioned studies. As described above, we observed significant enrichment between the MAL^MSC^ group and the TCGA MITF-low signature (Extended Data Fig. 8d). We also observed significant enrichment between the MAL^ADT^ group and the Cirenajwis *et al*. “normal-like” signature, consistent with these cells retaining a substantial component of the differentiated melanocyte program. Surprisingly, neither the MAL^FET^ nor the MAL^NEO^ group segregated with previously defined signatures. The MAL^FET^ group was significantly enriched for both the TCGA immune and MITF-low signatures, whereas the MAL^NEO^ group was not enriched for any previously defined signature. These observations suggest that categorization of malignant melanoma cells by the human developmental stage categories defined here represents a novel classification system, with the MAL^MSC^ and MAL^ADT^ groups reasonably aligned with previously reported MITF-low/stem cell and normal melanocytes, respectively, and the MAL^FET^ and MAL^NEO^ groups representing previously unreported signatures. While there was no significant enrichment for the *in vitro* differentiation based gene signatures from Tsoi *et al*. in any of the MAL groups, our analyses suggests that sequential dedifferentiation, which recapitulates the ordered cascade of differentiation in reverse, is predominant in melanoma progression (Extended Data Fig. 8a). This discovery mirrors the findings of Tsoi *et al*. which show that development of therapeutic resistance in melanoma traverses a sequential dedifferentiation trajectory^19^.

### SUPPLEMENTARY DATA

**Supplementary Table 1:** Related to Fig1 and Fig 2. Donor demographics and cell counts for skin samples used for single cell RNAseq.

**Supplementary Table 2:** Related to Fig. 2. Differentially expressed genes between donor-matched volar and non-volar cutaneous melanocytes.

**Supplementary Table 3:** Related to Extended Data Fig. 5. Pairwise differentially expressed genes between melanocyte developmental stages.

**Supplementary Table 4:** DevMel profile genes from Fig. 3.

**Supplementary Table 5:** Melanoma dedifferentiation analysis gene set from Fig. 3

**Supplementary Table 6:** SKCM group categorization of TCGA tumors in Fig. 4 and list of immune and FDA drug targets in Extended Data Fig. 8

### METHODS

#### Human Subject details

All skin was collected from surgical discards with informed consent and approval from the UCSF Institutional Review Board. All ages, races/ethnicities, and sexes were included in the eligibility criteria for this study. Adult tissue was obtained from surgical remnants of heathy skin taken for reconstructive surgery or from amputations with heathy skin. Neonatal foreskins were obtained after routine circumcision. Anonymous fetal specimens were obtained from elective terminations and fetal age (stated as fetal weeks) was estimated by heel-toe length^37^. When possible, fetal sex was determined by visual inspection using a dissecting microscope. All samples were collected in cold CO2 Independent Media (Gibco–Thermo Fisher Scientific) or Medium 154 (Gibco) with 1× Antibiototic-Antimycotic (Gibco) at 4°C until dissociation.

#### Skin sample preparation

Tissue dissociation was started the same day as sample acquisition. For adult and neonatal skin, the epidermis was enzymatically dissociated from the dermis with a dispase, neutral protease, grade II (Roche–Sigma-Aldrich), incubation for 14 hours at 4°C. Epidermal sheets were manually separated from the dermis, finely minced, and incubated with 0.5% trypsin (Gibco) for 3 minutes at 37°C. After manual trituration, trypsin was deactivated using ice cold soybean trypsin inhibitor (Gibco), then diluted 2:3 in ice cold Hanks’ balanced salt solution, no Mg2+, no Ca2+ (Gibco). The dissociated cell suspension was centrifuged at 500g, 4°C, for 4 minutes, resuspended in FACS buffer (0.1% bovine serum albumin (Sigma) and 25mM Hepes (Gibco) in Dulbecco’s phosphate-buffered saline (DPBS) (Gibco)) and strained with a 70μM filter to achieve a single cell suspension. For fetal tissue, the developing epidermis was manually removed from the dermis following a 20 – 30 minute incubation with 10mM EDTA (Invitrogen), DPBS at 37°C. The resulting epidermal layer was incubated with 0.5% trypsin (Gibco) for 1 min at 37°C and manually triturated. Trypsin was deactivated using ice cold soybean trypsin inhibitor (Gibco), then diluted 2:3 in ice cold Hanks’ balanced salt solution (Gibco). The dissociated cell suspension was centrifuged at 500g, 4°C, for 4 minutes, resuspended in FACS buffer, and strained with a 70μM filter to achieve a single cell suspension.

#### FACS analysis and single cell sorting

Single cell suspensions were counted, diluted to 1×10^6^ cells/100ul with ice cold FACS buffer containing dye conjugated antibodies (anti-CKIT (104D2), 15ng/100μl (CD11705, Thermo Fisher Scientific), anti-ITGA6 (GoH3), 15ng/100μl (12-0495-82, Thermo Fisher Scientific) and CD11c, 1:20 dilution (46-0116-41, Thermo Fisher Scientific)) and incubated on ice for 25 minutes. Cells were washed one time with 10x volume of FACS buffer, centrifuged for 2 minutes at 500g, resuspended in 30ng/mL Dapi (D3571, Molecular Probes), FACS buffer. Resuspended cells were strained through a 35 μm nylon mesh filter and kept on ice until sorted.

Single cells were sorted into 384-well plates using the “Ultra purity” setting on a SH800S (Sony) sorter. For a typical sort, a tube containing 0.3-1ml the pre-stained cell suspension was vortexed gently and loaded onto the FACS machine. A small number of cells were flowed at low pressure to check cell concentration and amount of debris. Then the pressure was adjusted, flow was paused, the first destination plate was unsealed and loaded. Single cells were sorted into plates by gating to exclude dead/dying cells (DAPI+) and doublets. The majority of the plate contained melanocytes (CD11c-/CKIT+) with 4 to 5 columns of basal keratinocytes (CD11c-/CKIT-/ITGA6+) and other triple negative cells such as suprabasal keratinocytes (CD11c-/CKIT-/ITGA6-). Immediately, after sorting, plates were sealed with a pre-labeled aluminum seal, centrifuged at 4°C and flash frozen on dry ice, before storage at −80 for later use.

#### Lysis plate preparation

Lysis plates were created by dispensing 0.4 μl lysis buffer (0.5U Recombinant RNase Inhibitor (Takara Bio, 2313B), 0.0625% TritonTM X-100 (Sigma, 93443-100ML), 3.125 mM dNTP mix (Thermo Fisher, R0193), 3.125 μM Oligo-dT30VN (IDT, 5’AAGCAGTGGTATCAACGCAGAGTACT30VN-3’) and 1:600,000 ERCC RNA spike-in mix (Thermo Fisher, 4456740)) into 384-well hard-shell PCR plates (Biorad HSP3901) using a Tempest liquid handler (Formulatrix). All plates were then spun down for 1 minute at 3220xg and snap frozen on dry ice. Plates were stored at −80°C until used for sorting.

#### cDNA synthesis and library preparation

cDNA synthesis was performed using the Smart-seq2 protocol (Picelli et al., 2013, 2014). Briefly, 384-well plates containing single-cell lysates were thawed on ice followed by first strand synthesis. 0.6 μl of reaction mix (16.7 U/μl SMARTScribe Reverse Transcriptase (Takara Bio, 639538), 1.67 U/μl Recombinant RNase Inhibitor (Takara Bio, 2313B), 1.67X First-Strand Buffer (Takara Bio, 639538), 1.67 μM TSO (Exiqon, 5’-AAGCAGTGGTATCAACGCAGACTACATrGrG+G-3’), 8.33 mM DTT (Bioworld, 40420001-1), 1.67 M Betaine (Sigma, B0300-5VL), and 10 mM MgCl2 (Sigma, M1028-10X1ML)) was added to each well using a Tempest liquid handler or Mosquito (TTP Labtech). Reverse transcription was carried out by incubating wells on a ProFlex 2×384 thermal-cycler (Thermo Fisher) at 42°C for 90 min and stopped by heating at 70°C for 5 min. Subsequently, 1.5 μl of PCR mix (1.67X KAPA HiFi HotStart ReadyMix (Kapa Biosystems, KK2602), 0.17 μM IS PCR primer (IDT, 5’-AAGCAGTGGTATCAACGCAGAGT-3’), and 0.038 U/μl Lambda Exonuclease (NEB, M0262L)) was added to each well with a Mantis liquid handler (Formulatrix) or Mosquito, and second strand synthesis was performed on a ProFlex 2×384 thermal-cycler by using the following program: 1. 37°C for 30 minutes, 2. 95°C for 3 minutes, 3. 23 cycles of 98°C for 20 seconds, 67°C for 15 seconds, and 72°C for 4 minutes, and 4. 72°C for 5 minutes. The amplified product was diluted with a ratio of 1 part cDNA to 10 parts 10mM Tris-HCl (Thermo Fisher, 15568025). 0.6 μl of diluted product was transferred to a new 384-well plate using the Viaflow 384 channel pipette (Integra). Illumina sequencing libraries were prepared as described in (Darmanis et al., 2015). Briefly, tagmentation was carried out on double-stranded cDNA using the Nextera XT Library Sample Preparation kit (Illumina, FC-131-1096). Each well was mixed with 0.8 μl Nextera tagmentation DNA buffer (Illumina) and 0.4 μl Tn5 enzyme (Illumina), then incubated at 55°C for 10 min. The reaction was stopped by adding 0.4 μl “Neutralize Tagment Buffer” (Illumina) and spinning at room temperature in a centrifuge at 3220xg for 5 min. Indexing PCR reactions were performed by adding 0.4 μl of 5 μM i5 indexing primer, 0.4 μl of 5 μM i7 indexing primer, and 1.2 μl of Nextera NPM mix (Illumina). All reagents were dispensed with the Mantis or Mosquito liquid handlers. PCR amplification was carried out on a ProFlex 2×384 thermal cycler using the following program: 1. 72°C for 3 minutes, 2. 95°C for 30 seconds, 3. 12 cycles of 95°C for 10 seconds, 55°C for 30 seconds, and 72°C for 1 minute, and 4. 72°C for 5 minutes. Library pooling, quality control, and sequencing. Following library preparation, wells of each library plate were pooled using a Mosquito liquid handler. Pooling was followed by two purifications using 0.7x AMPure beads (Fisher, A63881). Library quality was assessed using capillary electrophoresis on a Fragment Analyzer (Agilent) or Tapestation (Agilent), and libraries were quantified by qPCR (Kapa Biosystems, KK4923) on a CFX96 Touch Real-Time PCR Detection System (Biorad). Plate pools were normalized to 2 nM and equal volumes from library plates were mixed together to make the sequencing sample pool.

#### Sequencing libraries from 384-well plates

Libraries were sequenced on the NextSeq or NovaSeq 6000 Sequencing System (Illumina) using 2 x 100bp paired-end reads and 2 x 8bp or 2 x 12bp index reads. NextSeq runs used high output kits, whereas NovaSeq runs used either a 200 or 300-cycle kit (Illumina, 20012860). PhiX control library was spiked in at 1%.

#### Single-cell transcriptomic processing and analysis

Single cell reads were mapped to the human reference hg38 containing ERCC sequences using STAR aligner^38^. HTSeq^39^ was used to create gene count tables. These count tables were compiled and processed using Scanpy^40^. Low-quality cells were filtered based on the following criteria: number of genes < 1,250 OR number of reads < 50,000. Each gene in the transcriptome exhibited read counts in at least 3 cells. Cells exhibiting > 2-fold higher number of genes than average were labeled as putative doublets and removed. Iterative Louvain clustering yielded cell type-specific clusters, which were annotated using published marker genes based on inter-cluster differential expression analysis (two-sided Mann Whitney U test, Benjamini-Hochberg FDR < 5%). Briefly, Louvain clustering was performed on the k-nearest neighbor graph in principle component space of scaled highly variable genes. Cells were visualized using 2-dimensional UMAP embeddings. Cell cycle status was inferred by the mean ranked expression of marker genes, referred to as the cell cycle program score^41^. Cells below the 95^th^-percentile of the cell cycle program score were labeled non-cycling; conversely, cells equal to or greater than 95^th^-percentile of the cell cycle program score were labeled cycling. In order to control for variance introduced by disproportionate populations of cycling cells across groups, non-cycling cells were considered for downstream analyses. Thus, anatomical site-specific analyses and human melanocyte differentiation programs analyses were conducted on non-cycling cells.

#### Percent v-mel and c-mel

Top-10 cutaneous and top-10 volar DEGs were identified from the site-enriched genes based on highest median per-patient log-fold-change between cutaneous and volar samples. Individual cells were classified as v-mel if 4 or more top-10 volar DEGs exhibited non-zero expression AND fewer than 4 top-10 cutaneous DEGs exhibited non-zero expression. Conversely, individual cells were classified as c-mel if 4 or more top-10 cutaneous DEGs exhibited non-zero expression AND fewer than 4 top-10 volar DEGs exhibited non-zero expression. Percent v-mel and c-mel were then calculated for each skin specimen of unique anatomical location from each individual patient.

To determine the percent of HPGD positive melanocytes in tissue sections, melanocytes (TYPR1+ cells) were manually counted. Fraction of cells was determined by the number of HPGD+ TYRP1+ cells divided by the total number of TYRP1+ cells from each fixed frozen section. To quantify the number of NTRK2 and HPGD foci per DCT+ cells from the RNAscope data, images were processed to correct for Opal 570 (HPGD) bleed-through into the Opal 620 (NTRK2) channel. After bleed-through correction, DCT and associated dapi signal was used to define the area of DCT+ cells. NTRK2 and HPGD foci within DCT+ cells were then manually counted manually. All Image analysis was performed in Fiji (http://fiji.sc/)

#### Immunofluorescence

Skin samples were fixed in 4% paraformaldehyde (Electron Microscopy Sciences) at 4 °C overnight, washed with cold DPBS prior to paraffin or OCT embedding. Fixed frozen skin sections were incubated in blocking buffer: 2.5% donkey serum, 2.5% goat serum (Jackson ImmunoResearch Laboratories), 1% bovine serum albumin (Sigma-Aldrich), and 0.1% Triton X-100 (Sigma-Aldrich) for 1–2 hours at room temperature. The following primary antibodies were used at the indicated concentration in blocking buffer overnight at 4°C: mouse monoclonal anti-TYRP1 1:200 (TA99, ab3312, Abcam), mouse monoclonal anti-CKIT 1:100 (CD11700, Invitrogen–Thermo Fisher Scientific), rabbit polyclonal anti-HPGD 1:100 (HPA005679, Sigma-Aldrich). Secondary antibodies against mouse IgG, or rabbit IgG conjugated to DyLight 488 or 594 (Thermo Fisher Scientific) were used at a 1:1,000 dilution for 1–2 h at room temperature followed by Dapi, 1:1000 (Molecular Probes) for 1 minute. Sections were mounted in VECTASHIELD Vibrance (Vector Laboratories) prior to imaging.

Immunofluoresence images were acquired using Nikon NIS-Elements multi-platform acquisition software on an fully automated Nikon Ti-E inverted microscope with an Apo TIRF, 60x, 1.49 NA, oil objective (Nikon) and a Clara CCD camera (Andor).

#### Multiplex RNA-FISH

We performed Muliplex RNA-FISh using the RNAscope Assay RNAscope Multiplex Fluorescent V2 assay (Bio-techne, Cat. No. 323110) kit according to manufacturer’s protocol on 10 μM FFPE tissue sections. Tissues were stained using probes purchased from ACD for *HPGD* (Channel 1), *NTRK2* (Channel 2), *DCT* (Channel 3) and TSA Opal 570 (Channel 1, Akoya Biosciences, Cat. No. FP1488001KT), TSA Opal 620 (Channel 2, Akoya Biosciences, Cat. No. FP1488001KT) and TSA Opal 690 (Channel 3, Akoya Biosciences, Cat. No. FP1497001KT). TSA was used at a 1:1500 dilution. Cells were counterstained with DAPI and mounted with Prolong Gold Antifade Mountant (Thermo Fisher, P36930). Tissues were imaged using an Leica DMi8 microscope.

#### Diffusion Pseudotime

Diffusion pseudotime analysis on all non-cycling melanocyte cells was performed using the ‘scanpy.tl.dpt’ function. The pseudotime reference root cell was chosen from the youngest sample (9.5 f.w.). The diffusion map was computed from an n=30 neighborhood graph with a Gaussian kernel.

#### Single cell developmental stage melanonocyte (DevMel) logistic regression model

Input data was composed of single cell transcriptomes from the following 4 groups: MSC, FET, NEO and ADT. The input examples were randomly sampled and the number of examples was balanced among all labels. The combination of normal and melanoma transcriptomes was used to scale and center the data. The input data was split into testing and training partitions at a ratio 33:67. We implemented elasticnet regularization with an l1 ratio = 0.8. Single cell transcriptomes were evaluated by the model to yield a developmental stage label.

#### Classification of genes in melanoma dedifferentiation categories

Logistic regression variables and top-100 differentially expressed genes for each melanoma cell population grouped by DevMel label were used in subsequent dedifferentiation pathway analysis. DevMel group mean ranked expression was compared between normal and melanoma datasets to determine pathway based on the following criteria:

<u>Direct dedifferentiation</u>: All cancer DevMel group >= mean normal non-ADT DevMel group, 4-fold normal ADT DevMel group < mean normal non-ADT DevMel group
<u>Sequential dedifferentiation</u>: Max cancer DevMel group == Max normal DevMel group
<u>Melanoma-specific</u>: For each, cancer DevMel group > corresponding normal DevMel group, All cancer DevMel groups > 40^th^-percentile of expression, All normal DevMel groups < 10^th^-percentile of expression
<u>Normal-specific</u>: For each, cancer DevMel group < corresponding normal DevMel group, All cancer DevMel groups < 10^th^-percentile of expression, All normal DevMel groups > 15^th^-percentile of expression
  <u>Down regulated</u>: All cancer DevMel group <= mean normal non-ADT DevMel group, Normal ADT DevMel group > 1.5-fold mean normal non-ADT DevMel group
  <u>Not-readopted</u>: All cancer DevMel group <= normal ADT DevMel group, 1.5-fold normal ADT DevMel group < mean normal non-ADT DevMel group

#### Bulk tumor deconvolution

CIBERSORT^29^ was used to deconvolve bulk RNA-seq from the SKCM-TCGA cohort. As input, CIBERSORT requires cell type-labeled transcriptomes to estimate the proportion of each cell type in a bulk RNA-seq sample. Here, trimmed both single cell and bulk RNA-seq transcriptomes to include only genes that are shared in both datasets. Adopting a k-fold cross-validation approach, we prepared 10 sets of single cell input transcriptomes from normal melanocytes across 4 developmental stages: MSC, FET, NEO and ADT (balanced cell counts across all labels). Each input transcriptome set was used to devolve the SKCM-TCGA bulk RNA-seq samples, yielding 10 estimates of cell proportion. For each individual sample in the SKCM-TCGA dataset, the label means were used as the final estimate of label proportion. Hierarchical clustering was used to group SKCM-TCGA samples based on similar label proportions. One-sided Fisher Exact test was used to determine significant enrichment between two gene lists. The lifelines python package (10.5281/zenodo.3833188) was used to create Kaplan-Meier survival plots and perform logrank tests using curated SKCM-TCGA metadata^42^.

#### Data availability

Jupyter notebooks with detailed analysis scripts are available at: github.com/czbiohub/human_melanocytes

GEO: GSE151091

Bioproject: PRJNA625154

SAMN14593853 : A1015LM

SAMN14593854 : 12WKM01

SAMN14593855 : FS043_LM

SAMN14593856 : A1011L

SAMN14593857 : A1020LM

SAMN14593858 : A1033M

SAMN14593859 : A1022M

SAMN14593860 : A1014L

SAMN14593861 : A1038LM

SAMN14593862 : A1026L

SAMN14593863 : A1021M

SAMN14593864 : FS030_LM

SAMN14593865 : A1016LM

SAMN14593866 : A1025L

SAMN14593867 : A1017LM

SAMN14593868 : A1012M

SAMN14593869 : 18WKM06

SAMN14593870 : 12WK05

SAMN14593871 : 10WK03

SAMN14593872 : 16WKM04

SAMN14593873 : 9.5WK02

SAMN14593874 : A1046M

