## Supplementary material for "Human melanocyte development and melanoma dedifferentiation at single cell resolution": Table S1

**Supplementary Table 1: Donor demographics and cell counts for skin samples used for single cell RNAseq.**

| sample | age^+^ | sex | Anatomical location | skin  Pigmentation level ^++^ | Race / ethnicity | total # of melanocytes^+++^ |
| --- | --- | --- | --- | --- | --- | --- |
| 9.5WK02 | 9.5 f.w. | unknown | leg  arm | n/a  n/a | unknown | 145 |
| *10WK03 | 10 f.w. | unknown | leg  arm  palm | n/a  n/a  n/a | unknown | 759 |
| *12WK01 | 12 f.w. | M | leg  arm  palm  sole | n/a  n/a  n/a  n/a | unknown | 625 |
| *12WK05 | 12 f.w. | unknown | leg  arm  sole | n/a  n/a  n/a | unknown | 552 |
| 16WK04 | 16 f.w. | unknown | palm | n/a | unknown | 18 |
| *18WK06 | 18 f.w. | unknown | leg  palm  sole | n/a  n/a  n/a | unknown | 332 |
| FS030 | 0 yr | M | foreskin | LM | unknown | 251 |
| FS043 | 0 yr | M | foreskin | LM | unknown | 501 |
| a1021 | 24 yr | F | leg | M | Asian | 233 |
| *a1038 | 35 yr | M | leg  sole | LM  L | White | 610 |
| a1012 | 37 yr | F | arm | M | Asian | 82 |
| a1022 | 42 yr | M | arm | M | Hispanic or Latino | 311 |
| a1015 | 52 yr | F | arm | LM | Asian | 26 |
| a1025 | 56 yr | M | leg | L | White | 230 |
| a1016 | 58 yr | M | arm | LM | White | 225 |
| a1020 | 60 yr | F | arm | LM | Asian | 233 |
| a1033 | 61 yr | M | leg | M | Hispanic or Latino | 236 |
| a1011 | 65 yr | M | arm | L | White | 360 |
| a1026 | 66 yr | M | leg | L | White | 217 |
| a1014 | 68 yr | F | leg | L | White | 216 |
| ­ | 77 yr | F | leg  sole | M  L | Hispanic or Latino | 871 |
| a1017 | 81 yr | M | leg | LM | Hispanic or Latino | 55 |

^*^ included in multi-site patient matched analysis

^+^ f.w., fetal weeks; yr, years

^++^ pigment score based on appearance of skin prior to epidermal dissociation: L, light; LM, light-medium; M, medium

^+++^ number of melanocytes that passed quality control
